# Editing strigolactone biosynthesis genes in tomato reveals novel phenotypic effects and highlights *D27* as a key target for parasitic weed resistance

**DOI:** 10.1101/2025.02.01.636032

**Authors:** A. Nicoliaa, A. Cuccurullo, K. Tamada, K. Yoneyama, J.L. Rambla, A. Granell, F. Camerlengo, G. Festa, G. Francese, F. Contaldi, A. D’Alessandro, M.M. Rigano, L. Principio, N. D’Agostino, T. Cardi

**Affiliations:** CREA, Research Centre for Vegetable and Ornamental Crops, Pontecagnano (SA), Italy; Department of Agricultural Sciences, University of Naples Federico II, Portici (NA), Italy; Graduate School of Agriculture, Ehime University, Ehime, Japan; Department of Biochemistry and Molecular Biology, Saitama University, Saitama, Japan; Department of Biology, Biochemistry and Natural Sciences, Jaume I University, Castellón de la Plana, Spain; Instituto de Biología Molecular y Celular de Plantas (CSIC-UPV), Valencia, Spain; CNR-IBBR, Institute of Biosciences and Bioresources, Portici (NA), Italy

**Keywords:** Biotic stress, broomrapes, Orobanche, Phelipanche spp., genome editing, phenotyping, fruit-related traits, volatilome

## Abstract

Parasitic weed infestations pose an increasing threat to agriculture worldwide, especially in the Mediterranean region. *Phelipanche ramosa* and *P. aegyptiaca* (broomrapes) cause severe damage to field-grown tomato (*Solanum lycopersicum L.*). Strigolactones (SLs), apocarotenoid phytohormones, play a critical role in plant physiology and development, and are also the primary signals that trigger the germination of parasitic weed seeds. We generated CRISPR/Cas9 tomato knock-out lines for the *SlD27* gene, as well as three other key genes involved in SL biosynthesis (*SlCCD7*, *SlCCD8*, *SlMAX1*), all within the same genetic background. The edited lines exhibited a marked reduction in SL content in root exudates, along with impaired broomrape seed germination. A comprehensive analysis of morphological, reproductive, and fruit-related traits revealed gene-specific effects on plant phenotype, including vegetative traits, fruit set, fruit development, and volatilome. Specifically, the knock-out of two *CCDs* and the *MAX1* had a specific impact not only on plant development but also on the production of volatile organic compounds during fruit ripening. In contrast, the *Sld27* lines, produced for the first time in this study, displayed a phenotype similar to the control non-edited plants, suggesting that the *D27* gene holds promise as a breeding target for enhancing resistance to parasitic weeds in tomato.

**Highlight:** The characterization of tomato CRISPR/Cas9-edited lines for the four core genes involved in strigolactone biosynthesis revealed gene-specific effects on plant phenotype, with *D27* emerging as a potential target for resistance to parasitic weeds.

## Introduction

Parasitic weeds are plants that have partially (hemiparasites) or entirely (holoparasites) lost their autotrophic ability and depend on external hosts for water and nutrients (Vurro *et al*., 2019). Broomrapes (*Orobanche* and *Phelipanche spp*.), holoparasites of the *Orobanchaceae* family, establish parasitism by forming a vascular connection with host roots and are significant agricultural pests in the Mediterranean region (Cuccurullo *et al*., 2022). Among vegetable crops, tomato (*Solanum lycopersicum* L.) is particularly affected by *Phelipanche ramosa* L. and *P. aegyptiaca* L., leading to substantial yield losses (Vurro *et al*., 2019). Identifying novel tomato varieties/genotypes resistant to parasitic weeds is therefore highly desirable. The use of new genomic techniques (NGT), such as genome editing, could accelerate the development of resistant varieties in breeding programs (Yıldırım *et al*., 2024).

Strigolactones (SLs) are carotenoid-derived phytohormones exuded by plant roots to recruit arbuscular mycorrhiza fungi (AMF) under nutrient-deficient conditions (*e.g.* phosphorous and nitrogen). Furthermore, they serve as the primary signal inducing the germination of parasitic weed seeds (Mashiguchi *et al*., 2021).

Altering SL biosynthesis, and thereby reducing their release into the rhizosphere, prevents the germination of parasitic weed seeds in the soil, disrupting the initial phase of infection (*i.e.* pre-attachment) (Fernández-Aparicio *et al*., 2016). Extensive research on SL biosynthesis has been conducted in model species (*e.g. Arabidopsis thaliana* L.) and selected crops (*e.g.* rice, tomato; reviewed in Al-Babili and Bouwmeester, 2015; Mashiguchi *et al*., 2021; Dun *et al*., 2023).

In tomato, SLs are synthesized through a series of sequential reactions primarily localized in the plastid. The conversion of all-trans-β-carotene to carlactone is catalyzed by a carotenoid isomerase (Dwarf27 – D27) and two Carotenoid Cleavage Deoxigeneases (CCD7 and CCD8) (Vogel *et al*., 2010; Kohlen *et al*., 2012). Carlactone is further oxidated to carlactonic acid (CLA) by the cytosolic enzyme More Axillary Growth (MAX1) (Zhang *et al*., 2018). Subsequent oxidation steps, catalyzed by various cytochrome P450 enzymes, produce canonical SLs, including orobanchol, didehydro-orobanchol and solanacol (Wakabayashi *et al*., 2019; Wang *et al*., 2022).

Tomato lines altered in the function of core SL pathway genes (*CCD7*, *CCD8* and *MAX1*) have been previously developed demonstrating reduced colonization by parasitic weeds and providing valuable insight into potential future solutions for mitigating the impact of such parasitic weeds on tomato (reviewed in Cuccurullo *et al*., 2022). They also exhibited pleiotropic effects, including increased shoot branching and reduced height, confirming the multiple roles of SLs in plant development and physiology (Mashiguchi *et al*., 2021). However, previous studies on the genes involved in SL biosynthesis were conducted in various genetic backgrounds (reviewed in Cuccurullo *et al*., 2022), complicating direct comparisons of the observed phenotypes. Moreover, specific functional studies on the carotenoid isomerase D27 in tomato and other crops, as well as its relevance in conferring resistance to parasitic weeds, remain unexplored.

In this study, we generated CRISPR/Cas9 tomato knock-out (KO) lines for the *SlD27* gene and three other key SL biosynthetic genes (*SlCCD7*, *SlCCD8*, *SlMAX1*) within the same genetic background. We evaluated the SL content in root exudates and assessed the germination rates of *P. ramosa* and *P. aegyptiaca* in an *in vitro* assay. Furthermore, to compare the effects of inactivating the different genes, and assess the breeding and agronomic value of the edited lines, we performed a comprehensive characterization of morphological, reproductive and fruit-related traits, including the characterization of the volatilome.

## Materials and methods

A detailed description of the materials and methods used in this work can be found in the supplementary materials.

### Generation of CRISPR/Cas9 knock-out plants

The design of the single-guide RNAs (sgRNA) and the prediction of off-targets for the Sl*D27*, Sl*CCD7*, Sl*CCD8* and Sl*MAX1* genes were performed with the software CRISPR-P 2.0 (Liu *et al*., 2017). CRISPR/Cas9 vector assembly was performed via the Golden Gate Cloning method (Engler *et al*., 2014). Vector efficiency was preliminary assessed in tomato hairy roots (HRs) (Ron *et al*., 2014). The four validated vectors were independently introduced into *A. tumefaciens* strain EHA105. Transformation of tomato cotyledons (*S. lycopersicum* L. cv. *Ailsa Craig*) was carried out following the protocol by Qiu *et al*. (2007). At least two independent knock-out (KO) lines per gene, each showing an out-of-frame insertion in the coding sequence, were selected and advanced to the T_2_ generation through self-pollination. All the PCR primers used are listed in the Supp. Table S1.

### Parasitic weed germination assay

Tomato plants were grown in pots (3 to 4 biological replicates per genotype) filled with vermiculites under a 16 h light (approximately 240 µmol m^−2^ s^−1^) and 8 h dark cycle at 23-25°C. To collect root exudates containing SLs, tap water (100-200 mL) was poured onto the soil surface, and the drained solutions containing root exudates were collected. These solutions were extracted with ethyl acetate, and the ethyl acetate phase was subsequently dried over anhydrous MgSO_4_ and concentrated *in vacuo*. All crude samples were stored at 4°C until use. These root exudate samples were used for both germination assay and LC-MS/MS analysis. Germination assays with broomrape seeds were conducted as reported previously (Yoneyama *et al*., 2007). Seeds germination was assessed by counting seeds where the radicle had protruded through the seed coat.

### LC-MS/MS analysis of root exudates

Tomato SLs were analyzed using ultra-performance liquid chromatography coupled with tandem mass spectrometry (UPLC-MS/MS) following the protocol described by Yoneyama *et al*., (2022). The analysis was performed on a Acquity UPLC System (Waters, Milford, Massachusetts, USA) connected to a Xevo TQD triple-quadrupole mass spectrometer (Waters Milford, Massachusetts, USA) with an electrospray ionization (ESI) interface. Multiple reaction monitoring (MRM) transitions were used to detect specific SLs.

### Detection of plant morphological, reproductive and fruit-related traits

Four biological replicates for each of the T_2_ tomato biallelic homozygous edited lines *Sld27* (*d27*-KO1, *d27*-KO2), *Slccd7* (*ccd7*-KO1, *ccd7*-KO2, *ccd7*-KO3), *Slccd8* (*ccd8*-KO1, *ccd8*-KO2, *ccd8*-KO3) and *Slmax1* (*max1*-KO1, *max1*-KO2), as well as non-edited control plants, were transplanted into pots containing a mixture of peat and perlite. These plants were grown in a phytotron under regular watering and fertilizing conditions, with the pots arranged in a randomized design. The morphological, reproductive and fruit-related traits were recorded at specific days after transplanting (DAT). Ten fruits per biological replicate were tagged at the breaker stage. Fruits were harvested after 7 days (breaker + 7) measured and subsequently used for biochemical analysis. Five additional fruits per biological replicate were harvested at the mature green stage and stored in a controlled environment to estimate water loss. The number of adventitious roots was counted on plants at 120 DAT.

### Extraction and U/HPLC-PDA analysis of chlorogenic acid, rutin, naringenin and sugars

Each biological replicate was analyzed in duplicate for simple sugars (fructose, glucose) for the two flavonoids naringenin chalcone (NaCh) and rutin (Rut), and the phenol chlorogenic acid (CA). The CA and the NaCh and Rut were extracted following a previously described method (de Vos *et al*., 2007), with minor modifications. The analyses were performed using an Ultimate 3000 UPLC system (Thermo Fisher Scientific, Sunnyvale, CA, USA.). The extraction of simple sugars was performed on 250 mg of lyophilized powder and extract subjected to analysis on the E-Alliance HPLC system (Waters, Milford, Massachusetts, USA), with data acquired and analyzed using Waters Empower software. Peak areas for CA, NACh, Rut and sugars were recorded using authentic, distinct, and appropriately diluted standards (Sigma-Aldrich, Saint Louis, MO, USA). Finally, compound content was expressed as fold change relative to the non-edited control plants.

### Extraction and HPLC-PDA analysis of carotenoids and tocopherols

Carotenoids and tocopherols were extracted following the protocols described by Barja *et al*. (2021) and Ezquerro *et al*. (2022) with modifications. Liquid chromatography was performed using an HPLC Water Alliance 2695 Separations module, with carotenoids detected by a Waters 996 PDA detector and tocopherols identified using a Waters 2475 Multi λ fluorescence detector. Peak areas were recorded and analyzed using Waters Millennium software (Waters, Milford, Massachusetts, USA). The areas were normalized by internal standard, and carotenoid and tocopherols content was expressed as a fold change relative to the non-edited control plants.

### Extraction and GC/MS analysis of volatile organic compounds

Volatile organic compounds (VOCs) were extracted and analyzed as reported in Rambla *et al*., (2017). For the analysis, 500 mg of fresh pericarp powder, obtained from the five pooled fruits per biological replicate and stored at −80°C, were used. VOCs were captured using headspace solid-phase microextraction (HS-SPME) and subsequently separated and identified by gas chromatography coupled with mass spectrometry (GC/MS). VOCs from the headspace were extracted using a 65 µm PDMS/DVB solid-phase microextraction fiber (SUPELCO-MERCK, Darmstadt, Germany). Desorption of the compounds from the fiber occurred at the injection port of a 6890N gas chromatograph (Agilent Technologies, Santa Clara, California, USA). Data acquisition was performed using a 5975B mass spectrometer (Agilent Technologies, Santa Clara, California, USA). Compound identification was accomplished by comparing retention times and mass spectra with those of pure standards. Peak areas were recorded and normalized results were expressed as the ratio of compound abundance in each sample relative to the reference admixture.

### Statistical analysis

Statistical analysis was performed using single factor analysis of variance (ANOVA) and Tukey’s HSD with the software package Real Statistic (https://real-statistics.com/). Statistical analysis on VOCs was performed by t-test with the software Office 365 Excel (Redmond, Washington, US). Linear Discriminant Analysis (LDA) was conducted using the MASS package in R (https://cran.r-project.org/web/packages/MASS/index.html).

## Results

### Knock-out mutants of strigolactone biosynthetic genes were efficiently generated using CRISPR/Cas9

The sgRNAs designed to specifically target genes involved in SL biosynthesis demonstrated substantial activity, with efficiencies ranging from 35 % to 89 %, in hairy root (HR) assays mediated by *Agrobacterium rhizogenes* (Supp. Table S2). Independent transformation of tomato cotyledons using *A. tumefaciens* and the same four vectors employed in the HR assays resulted in the generation of 10 KO plant lines: two lines each for *Sld27* and *Slmax1*, and three lines each for *Slccd7* and *Slccd8* (Fig. 1). Sanger sequencing of target genes from selected T_2_-generation plants (T-DNA free) confirmed that all edited lines were biallelic and homozygous for their respective mutations. Additionally, the presence of potential off-target effects was ruled out (Supp. Table S3). Editing in the newly generated *Sld27*, *Slccd7*, *Slccd8* and *Slmax1* lines resulted in frameshift mutations, leading to KO genotypes (Fig. 1). In the *Sld27* KO lines, the limited size of the exons hampered sgRNA design, necessitating targeting of the 5’ untranslated region. A single insertion was introduced before the start codon, accompanied by either a deletion (*Sld27*-KO1) or an insertion (*Sld27*-KO2) of a single base in the second exon (Fig.1). The three *Slccd7* KO lines exhibited deletions in the first exon: 1 bp in *Slccd7*-KO1 and *Slccd7*-KO2, and 2 bp in *Slccd7*-KO3, whereas the three *Slccd8* KO lines displayed a broader range of deletions in the second exon: 2 bp in *Slccd8*-KO2, 4 bp in *Slccd8*-KO1 and 5 bp in *Slccd8*-KO3. Finally, the two *Slmax1*-KO lines exhibited deletions in exon 3: 1 bp in *Slmax1*-KO2 and 4 bp in *Slmax1*-KO1 (Fig. 1)

**Figure 1.**
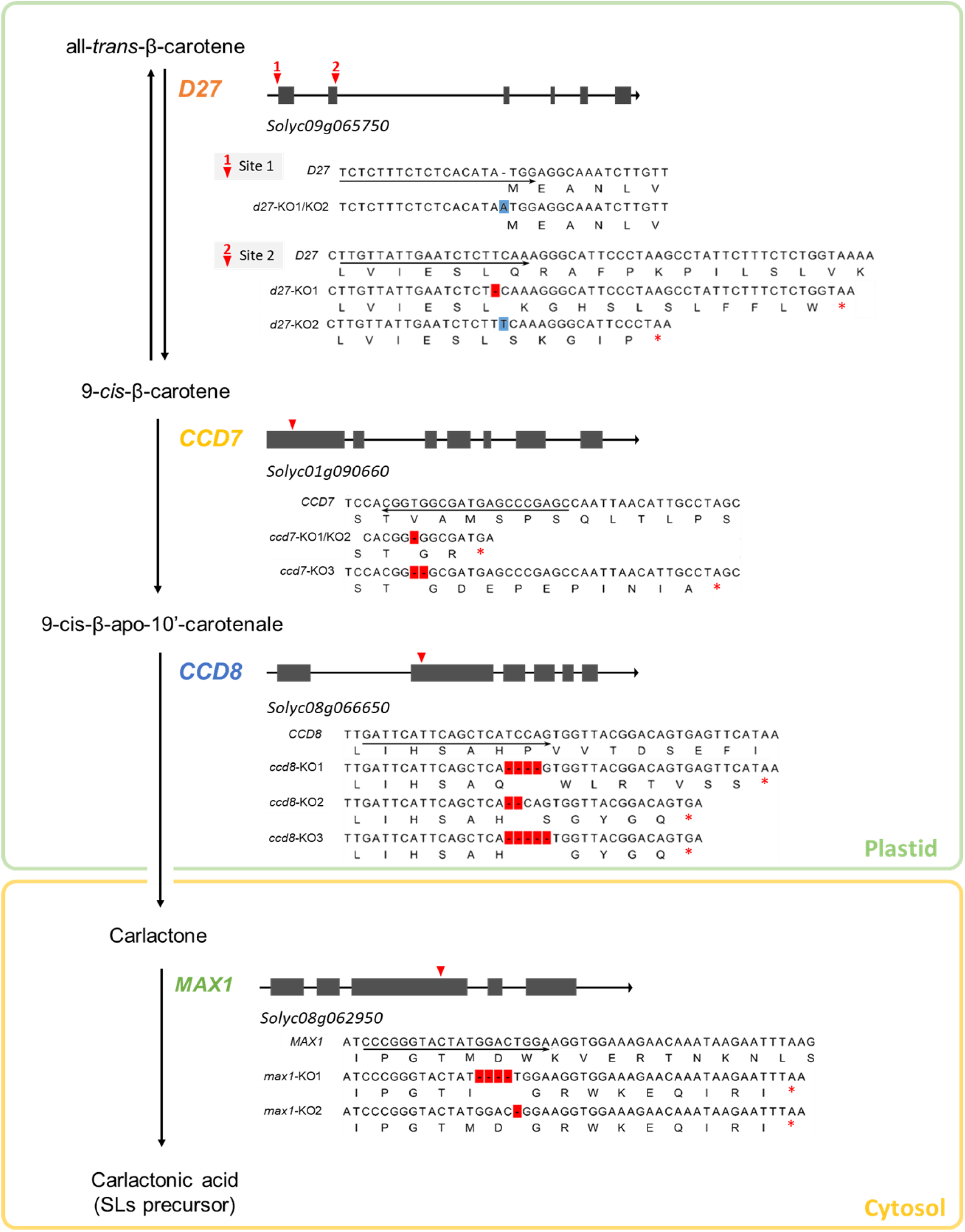
Schematic representation of induced mutations targeting core strigolactone biosynthesis genes in CRISPR/Cas9-edited tomato lines. The figure provides an overview of the compartmentalized strigolactone (SL) biosynthesis pathway in tomato, depicting the enzymatic steps from all-*trans*-β-carotene to carlactonic acid. The genomic structure of each SL biosynthesis gene is illustrated, with genes represented as black arrows oriented from the 5’(left) to the 3’(right) end (not drawn to scale). Exons are shown as grey boxes, and the CRISPR/Cas9 target sites are marked with inverted red triangles. Local alignment between the cDNA sequences of wild-type and CRISPR/Cas9 knock-out (KO) lines are displayed for each gene. The 20-nucleotide single guide RNA (sgRNA) sequence is underlined by a black arrow below the wild-type sequence, indicating its orientation. Mutations resulting from CRISPR/Cas9 activity, are highlighted in red (deletions) and blue (insertions) within the KO line sequences. Out-of-frame mutations are shown in the corresponding translated amino acid sequences below each cDNA sequence, with the premature stop codon indicated by a red asterisk.

### Strigolactones are not detectable in root exudates of edited tomato lines

Tomato plants produce and exude at least five SLs, including orobanchol, solanacol, 6,7-didehydroorobanchol, phelipanchol and epihelipanchol (Zhang *et al*., 2018; Wakabayashi *et al*., 2022). Authentic standards of orobanchol and solanacol were available, allowing for their confirmation. Both SLs were readily detectable in the root exudates of control non-edited plants but were absent in the edited tomato lines. This provided a functional confirmation that the four genes involved in the SL biosynthetic pathway were successfully knocked out by CRISPR/Cas9 (Supp. Fig. S1A, B). Authentic standards for 6,7-didehydroorobanchol, phelipanchol, and epihelipanchol were not available under our experimental conditions. Consequently, MRM channels for these SLs were set based on previously reported data (Zhang *et al*., 2018; Wakabayashi *et al*., 2022). A distinct single peak was observed exclusively in the root exudates of non-edited control plants and not in the edited lines (Supp Fig. S1C). However, it remained unclear whether this peak corresponded to 6,7-didehydroorobanchol, phelipanchol, or epihelipanchol.

### Root exudates from the edited tomato lines exhibited reduced germination activity for root parasitic weeds

We evaluated the effects of root exudates from non-edited control plants and CRISPR/Cas9 edited lines on the germination of *P. ramosa* and *P. aegyptiaca* (Fig. 2A, B), for which tomato is a regular host, and *Orobanche minor*, usually not found in tomato fields and used as non-host control (Supp. Fig. S2). The *Slccd7* and *Slccd8* lines were the most effective, inhibiting parasite germination by more than 90 %. A similar reduction was observed in the germination of *P. ramosa* seeds treated with *Sld27* root exudates. However, in *P. aegyptiaca*, residual germination rates up to approximately 21 % was observed with the *d27*-KO2 root exudates (Fig. 2B). Additionally, root exudates from *Slmax1* lines appeared to stimulate higher germination rates in *P. ramosa* and *O. minor* compared to *P. aegyptiaca* (Fig. 2A, B; Supp. Fig. S2)

**Figure 2.**
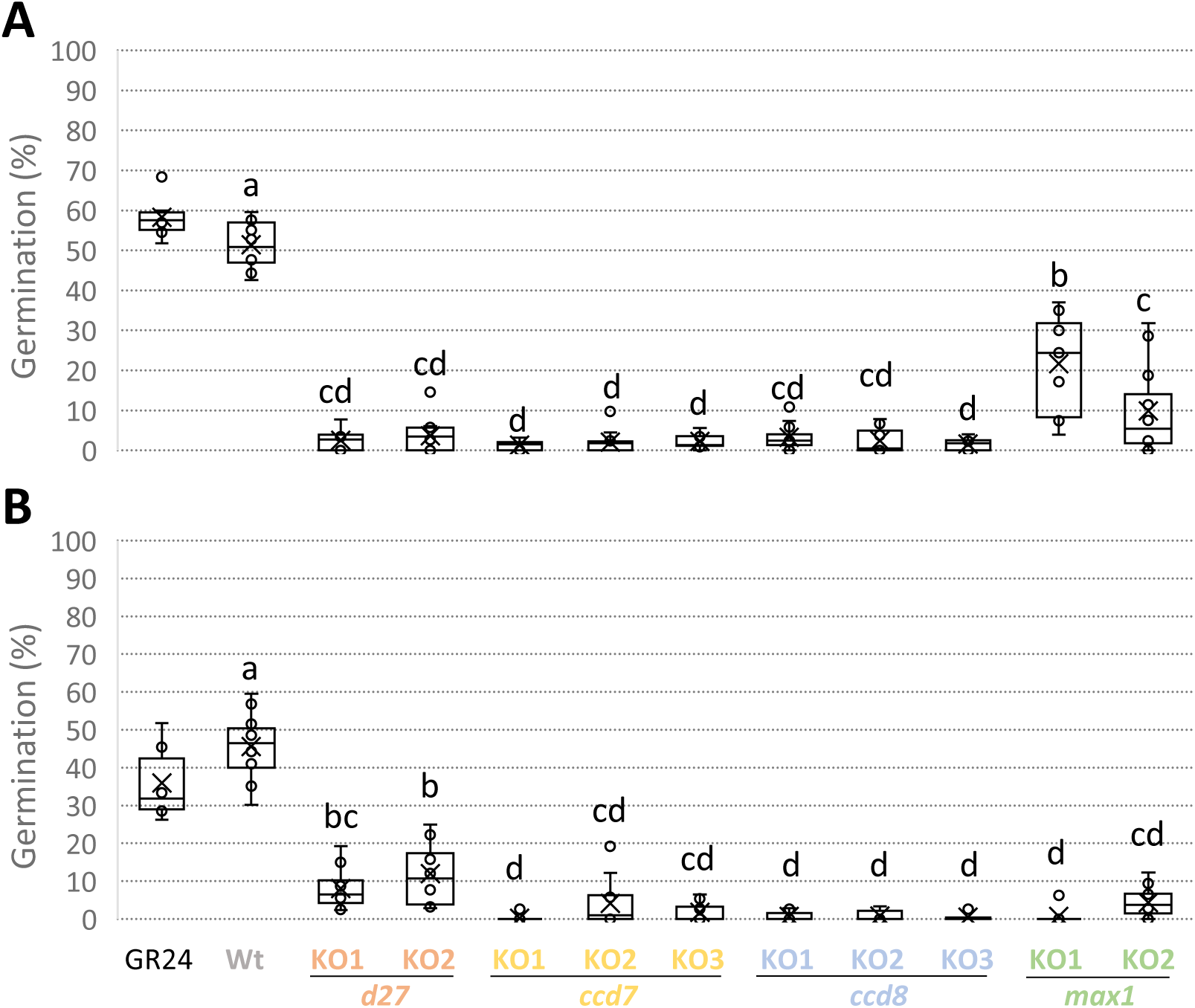
Germination-stimulating activity of tomato root exudates on *Phelipanche* spp. seeds. Root exudates were sampled from non-edited control plants (wild-type, Wt) and CRISPR/Cas9 knock-out (KO) lines targeting strigolactone biosynthesis genes (for each line, *n* = 3-4). These samples were used in germination assay with seeds of *Phelipanche ramosa* (A) and *P. aegyptiaca* (B). Statistically significant differences among genotypes are indicated by different letters (*p* < 0.05), as determined by Tukey’s HSD test. GR24 (10^-6^ M) was included as positive control in the assays.

### The Sld27 edited tomato lines exhibit morphological, reproductive, and fruit-related traits similar to those of the wild-type

The *Sld27*, *Slccd7, Slccd8* and *Slmax1* CRISPR/Cas9 KO lines generated in this study were compared with the control non-edited plants by evaluating a total of 13 phenotypic traits (Supp. Table S4).

Considering the morphological traits, height reduction was significant in *Slccd7*, *Slccd8* and *Slmax1* at all sampling points, due to a reduction in internode length particularly evident at 33 DAT. The increase in lateral branching was more pronounced in the *Slccd8* and *Slmax1* lines, which exhibited a doubling in the number of branches. The *Slccd7* and *Slccd8* lines showed a remarkable increase in adventitious root production compared to the control (up to 35-fold in *Slccd8*). Interestingly, the *Slmax1* lines displayed a smaller and less consistent increase in hairy roots (Supp. Table S4).

The analysis of reproductive traits evidenced that the *Slccd8* and *Slmax1* lines showed a significant reduction in flower numbers on the third inflorescence; notably, the *Slccd7* lines were comparable to the non-edited control plants. The *Slmax1* lines exhibited a remarkable reduction in fruit set (up to 73 %) a trait found also in the *ccd7*-KO3, whereas the *Slccd8* lines were similar to control (Supp. Table S4).

Interestingly, the examination of fruit-related traits highlighted an extended period from anthesis to the onset of ripening (up to 7 days) in the *Slmax1* lines. Aside from the marginal significance observed in *ccd7*-KO3, the other *Slccd7* lines, as well as the *Slccd8* lines, did not show significant changes.

The *Slmax1* lines exhibited also the most pronounced reduction in fruit weight (up to 17.58 grams). Similarly, two out of three *Slccd7* lines experienced a significant decrease (up to 11.34 grams), whereas among *Slccd8* lines, only *ccd8*-KO1 showed a significant reduction. Alterations in fruit width mirrored the changes in fruit weight, with the most severe reduction in *Slmax1* (up to 0.72 cm). Fruit length was also significantly reduced, with *Slmax1* lines showing the most pronounced reduction (up to 0.61 cm) (Supp. Table S4; Supp. Fig. S3).

The *Sld27* lines generally did not show significant differences in the morphological, reproductive and fruit-related traits compared to non-edited plants, evidencing a control-like phenotype (Supp. Table S4; Supp. Fig. S3).

These results were largely confirmed when the thirteen morpho-physiological traits were subjected to linear discriminant analysis (LDA) to verify the best separation of the lines according to the linear trait combinations and rank traits for their importance in group separation. The LDA model produced two canonical variables (*i.e.* discriminant functions), LD1 and LD2, which explained 79.08 % and 15.17 % of the variance, respectively (Supp. Table S5). Interestingly, the *Sld27* lines clustered very close to the non-edited control group (Fig. 3A) and were clearly distinguishable from the other lines (*Slccd7*, *Slccd8*, *Slmax1*), indicating a high degree of similarity between these mutants and the control in terms of the traits evaluated. LD1 was positively correlated (r ≥ 0.60) with plant height, average internode length, number of flowers on the third inflorescence, and fruit weight. Conversely, LD1 was negatively correlated (r ≤ −0.60) with the number of lateral branches, number of adventitious roots, and number of days between anthesis and the breaker stage. LD2 was positively correlated with the fruit set percentage and fruit width (Fig. 3B; Supp. Table S5).

**Figure 3.**
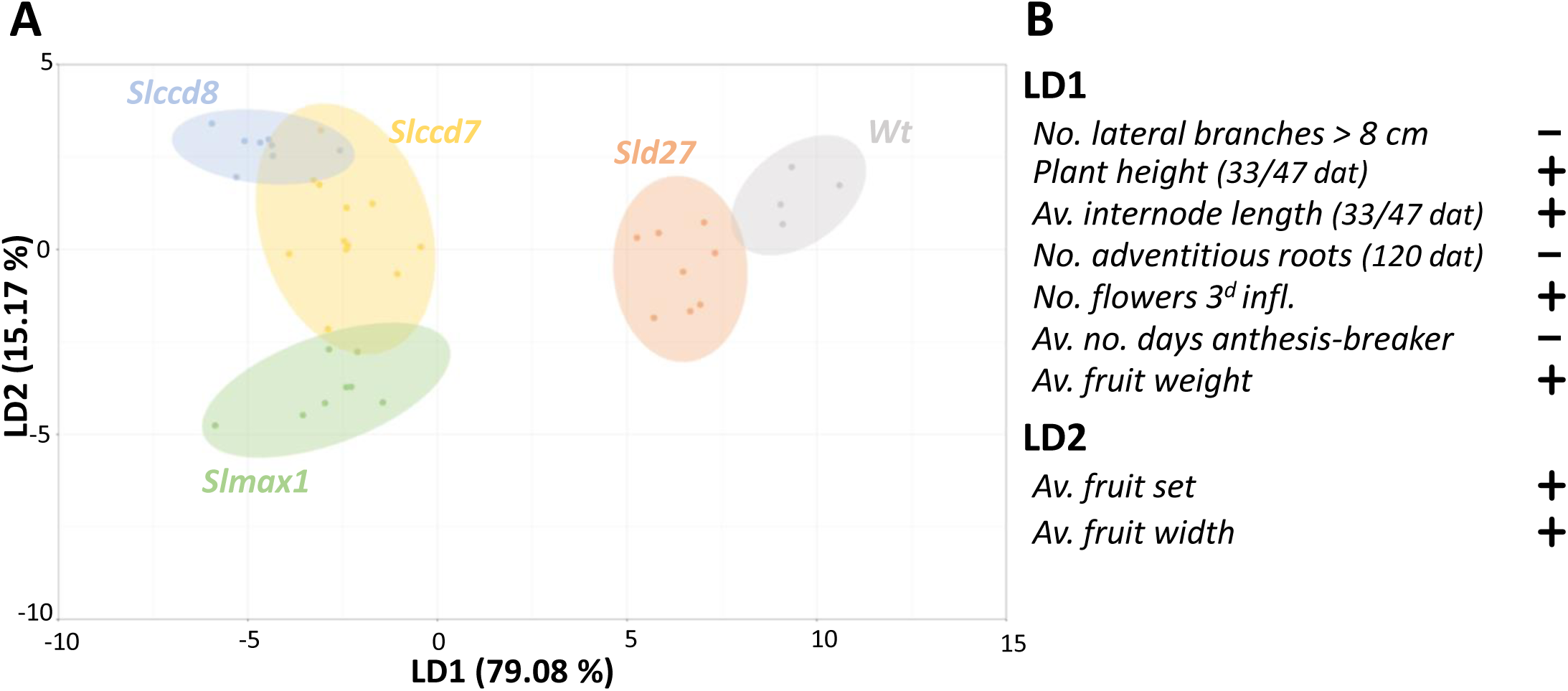
Bidimensional distribution of genotypes (ovals) and lines (dots) after linear discriminant analysis (LDA) of morphological, reproductive, and fruit-related traits. (A) Bidimensional distribution of scores assigned to CRISPR/Cas9 knock-out (KO) lines targeting strigolactone biosynthesis genes: *Sld27* (*d27*-KO1, *d27*-KO2; orange), *Slccd7* (*ccd7-*KO1, *ccd7-* KO2, *ccd7-*KO3; yellow), *Slccd8* (*ccd8-*KO1, *ccd8-*KO2; blue) and *Slmax1* (*max1-*KO1, *max1*-KO2; green). The non-edited control genotype (wild-type, Wt) is represented in grey (for each line, n=4). The two scores for each line were calculated using the linear discriminant functions LD1 (x-axis) and LD2 (y-axis), which together explain 94.25 % of the observed variability. (B) Traits with strong correlation to LD1 and LD2 are highlighted (+: positive corr., r ≥ 0.6; -: negative corr., r ≤ - 0.6). DAT: days after transplant.

### Profiling of fruit metabolites produced by edited tomato lines

The ANOVA and post-hoc analysis failed to clearly identify any specific line with differences in the average content of fructose and glucose between SL-depleted lines and the control (Supp. Table S6).

Regarding polar metabolites, quantification of chlorogenic acid showed no detectable level in the *ccd7*-KO3, whereas all other lines resembled the control (Supp. Table S7).

The analysis of the flavonoid rutin did not reveal statistically significant differences in post-hoc analysis between the edited lines and controls. Conversely, the flavonoid naringenin chalcone appeared to be generally increased, but statistically significant only in *max1*-KO1 (Sup. tab. S7).

Examining non-polar metabolites, the major carotenoids and tocopherols exhibited negligible changes, with the sole exception of the δ-tocopherol, that was reduced in *ccd8*-KO2 and the all-trans lycopene, which appeared slightly reduced in *max1*-KO2 (Supp. Table S8).

The ripe fruits unequivocally revealed sixty-four different VOCs. Specific VOC and metabolic pathways resulted highly affected by SL-depletion in tomato fruits (Supp. Table S9). Indeed, considering the benzenoid/phenylpropanoid pathway (B), eugenol, guaiacol and methyl salicylate were significantly increased (up to 2.02-, 2.8- and 8-fold, respectively) in the *Slmax1* lines. Among the branched-chain amino acids (BCAA) related compounds, 3-methyl butanoic acid was highly reduced in *Slccd7*, *Slccd8* and *Slmax1* (up to 0.29-fold). Similarly, 1-nitro-3-methylbutane was significantly reduced in both *Slmax1* lines (up to 0.12-fold). Within the phenylalanine derivative (Phe) group, the 2-phenylethanol was reduced in *Slccd8* and *Slmax1* lines (up to 0.29- and 0.16-fold, respectively). These declines were more pronounced for 1-nitro-2-phenylethane, which was also affected in the *Slccd7* lines. On the contrary, the pathways of fatty acid derivative (L), apocarotenoid (ApoC) and terpenoid (T) seemed to be not affected by mutations in SL genes (Supp. Table S9).

LDA was performed to summarize the behavior of the lines and confirm their separation based on VOC content, while simultaneously ranking the various compounds by their significance in group differentiation. Interestingly, the LDA identified four clusters: 1) non-edited control plants; 2) *Sld27* lines; 3) *Slccd7* and *Slccd8* lines; 4) *Slmax1* lines (Fig. 4A). The canonical variable LD1 accounted for 57.19% of the variability and was positively correlated (r ≥ 0.60) with two BCAA related compounds (3-methylbutanoic acid, 1-nitro-3-methylbutane), one L compound (E-2-pentenal) and two Phe compounds (2-phenylethanol, 1-nitro-2-phenylethane). Conversely, LD1 exhibited negative correlation (r ≤ −0.60) with two T compounds (p-cymene, 2-caren-10-al) (Supp. Table S10). The second canonical variable, LD2, explained 25.69% of the variability and was negatively correlated (r ≤ −0.60) with two B compounds (guaiacol, methyl salicylate) and positively correlated (r ≥ 0.60) with one BCAA-related compound (3-methylbutanenitrile) (Fig. 4B; Supp. Table S10).

**Figure 4.**
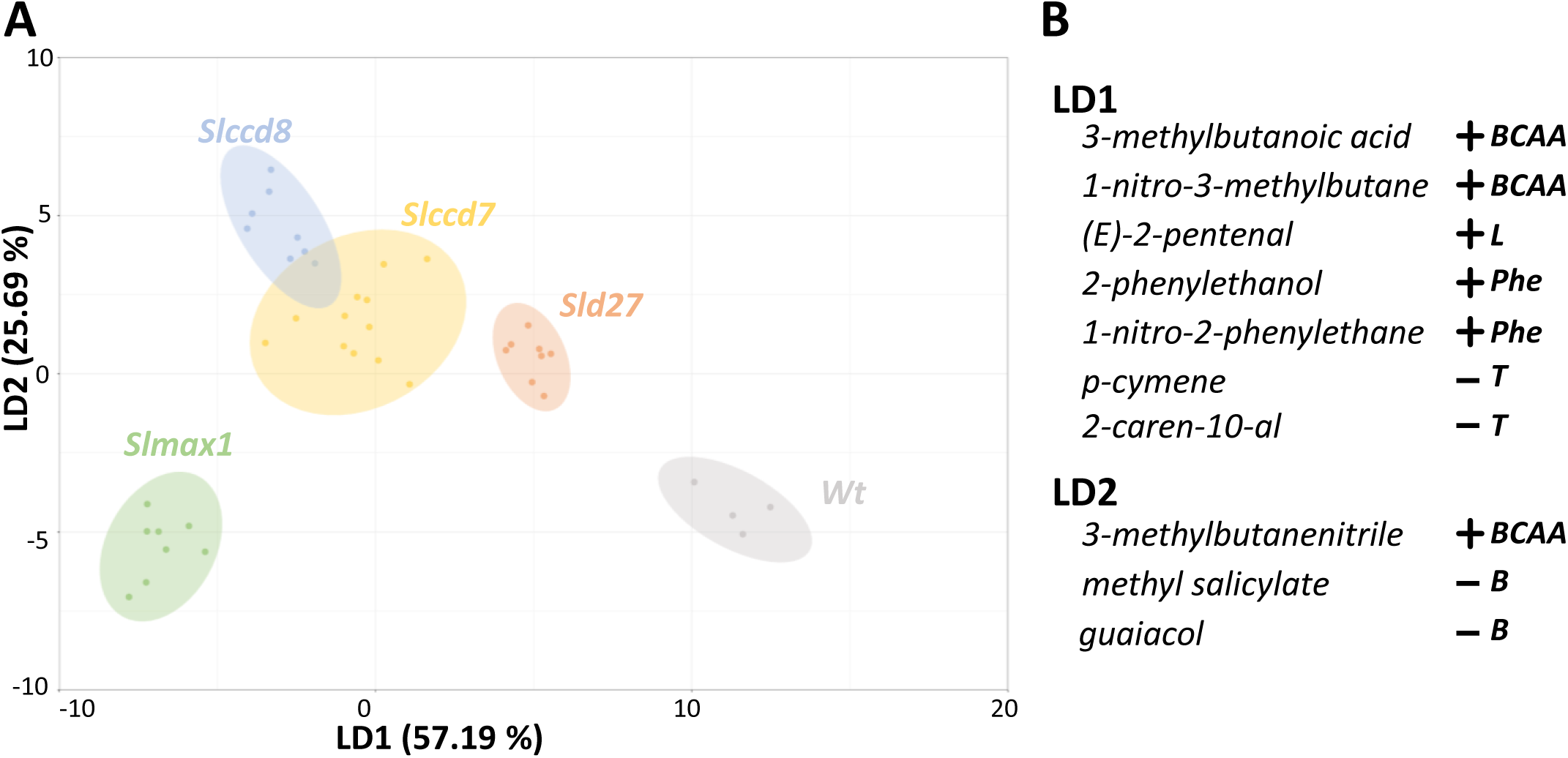
Bidimensional distribution of genotypes (ovals) and lines (dots) after linear discriminant analysis (LDA) of volatile organic compounds (VOCs) (A) Bidimensional distribution of scores assigned to CRISPR/Cas9 knock-out (KO) lines targeting strigolactone biosynthesis genes: *Sld27* (*d27*-KO1, *d27*-KO2; orange), *Slccd7* (*ccd7-*KO1, *ccd7-*KO2, *ccd7-*KO3; yellow), *Slccd8* (*ccd8-*KO1, *ccd8-*KO2; blue) and *Slmax1* (*max1-*KO1, *max1*-KO2; green). The non-edited control genotype (wild-type, Wt) is represented in grey (for each line, n = 4). Scores for each line were derived using linear discriminant functions LD1 (x-axis) and LD2 (y-axis), which together account for 82.88 % of the observed variability. (B) Selected VOCs with strong correlations to LD1 and LD2 are highlighted (+: positive corr., r ≥ 0.6; -: negative corr., r ≤ −0.6). Metabolic pathways associated with the VOCs are indicated as follows: BCAA, Branched-chain amino acid related; L, Fatty acid derivative; Phe, Phenylalanine derivative; T, Terpenoid; B, Benzenoid/Phenylpropanoid.

## Discussion

Various tomato lines with either complete or partial inactivation of genes involved in SL biosynthesis have been previously generated using different strategies and genetic backgrounds (Cuccurullo *et al*., 2022), thus preventing the direct comparisons of gene effects on parasitic plant resistance and on host plant phenotype. In this study, we successfully generated a complete panel of CRISPR/Cas9 KO lines targeting the core genes (*SlD27*, *SlCCD7*, *SlCCD8* and *SlMAX1*) involved in SL biosynthesis, all within the same cultivar, Ailsa Craig. Studying mutants in the same genetic background is valuable because it minimizes genetic variability and epistatic effects, allowing researchers to attribute observed phenotypic differences specifically to the mutation of interest. This approach ensures that differences in traits are not confounded by background genetic effects, leading to more accurate and reproducible conclusions. Additionally, using a consistent genetic background enhances the reliability of downstream analyses, and facilitates the identification of gene-specific roles in biological processes. This is particularly critical in complex traits like metabolism, development, or stress responses, where subtle genetic interactions can influence outcomes.

To assess the potential breeding value of these novel alleles as sources of resistance against parasitic weeds, we evaluated the biochemical composition of the root exudates and their effects on the germination of two common broomrapes species parasitizing tomato. Additionally, we performed a comprehensive phenotypic characterization of the edited lines, with a particular focus on the biochemical profile of the fruits.

### Phenotypic changes, varying in both intensity and specificity, are influenced by disruptions at different points along the SL biosynthetic pathway

Notably, this study represents the first characterization of a *d27* loss of function mutant in a dicotyledonous crop species. The absence of significant morphological changes observed in tomato *Sld27* lines, is consistent with the mild phenotypes reported in rice and Arabidopsis *d27* mutants (Lin *et al*., 2009; Waters *et al*., 2012). We have extended these findings to include reproductive and fruit-related traits in tomato. The *D27* gene is an iron-containing isomerase that is conserved across higher plants and algae but absent in animals and fungi (Waters *et al*., 2012). Previous studies on *D27* have been limited in scope, primarily focused on Arabidopsis, rice and the bryophyte *Marchantia polymorpha* (Lin *et al*., 2009; Waters *et al*., 2012; Jibran *et al*., 2024). Similar to rice and Arabidopsis, the *SlD27* gene is essential for SL biosynthesis in tomato, as evidenced by the absence of detectable SLs in root exudates of *Sld27* lines and the marked reduction in the germination observed in the parasitic plant species *P. ramosa* L. and *P. aegyptiaca* L. Similarly, root exudates from the Arabidopsis *d27* mutant caused a significant reduction in the germination rate of the parasitic weed *Striga hermonthica* L. (Yang *et al*., 2023). In rice *d27* mutant, the 2’-epi-5-deoxystrigolin was undetectable in the root exudates (Lin *et al*., 2009). The control-like phenotype observed in the tomato *Sld27* lines may be explained by non-enzymatic isomerization induced by light exposure, of all-trans-β-carotene to 9-cis-β-carotene (reviewed in Khoo *et al*., 2011), a phenomenon that could be sufficient to sustain SL biosynthesis in the epigeal part of the *Sld27* mutant. However, recent studies have demonstrated the existence of *D27* paralogues genes (*D27-Like1* and *D27-Like2*), in rice (Liu *et al*., 2020) and Arabidopsis (Gulyás *et al*., 2022; Yang *et al*., 2023). Notably, the *D27-Like1* isomerase has been shown to contribute, albeit to a lesser extent, to SL biosynthesis in Arabidopsis (Yang *et al*., 2023). In tomato, the presence of *D27-like* genes was predicted through phylogenetic analysis, although their importance in SL biosynthesis and plant development needs to be clarified (Cuccurullo *et al*., 2024).

In contrast to the *Sld27* lines, the *Slccd7*, *Slccd8* and *Slmax1* lines exhibited significant alterations in morphological, reproductive, and fruit-related traits.

The *Slccd*s lines were notably characterized by an increased production of adventitious roots compared to the other lines. This trait has been previously documented in *ccd7* and *ccd8* lines from various species (Rasmussen *et al*., 2012; Liu *et al*., 2013). For instance, Kohlen *et al*., (2012) reported an elevated number of adventitious roots in the stems of *Slccd8* RNAi-lines. This phenomenon is thought to be related to higher auxins concentrations and/or increased auxin sensitivity (Kohlen *et al*., 2012; Rasmussen *et al*., 2012). However, it remains intriguing to explore whether the extensive proliferation of hairy roots observed in the *Slccd7* and *Slccd8* lines is also linked to the absence of additional specific apocarotenoids produced by these two CCDs.

The *Slmax1* lines, edited in the downstream gene of SL biosynthetic pathway, exhibited unique phenotypes not previously evidenced or specifically associated (Zhang *et al*., 2018; Bari *et al*., 2021), with this genotype.

*Slmax1* lines retained a significant germination rate for *P. ramosa* and for the non-host species *O. minor*. The latter differs from *Phelipanche spp.* because it is not characterized by spontaneous germination, and it seems to be specifically sensitive to SLs and not to other germination inducers (*i.e.* isothiocyanates). Therefore, a possible explanation for this discrepancy may lie in the accumulation of carlactone in the *Slmax1* lines. In Arabidopsis, carlactone exhibits SL-like activity and has a clear stimulating effect on *S. hermonthica* seed germination (Alder *et al*., 2012). Although carlactone has been proposed as the possible mobile signal in plants (Al-Babili and Bouwmeester, 2015; Dun *et al*., 2023), it has not been detected in root exudates but only in root extracts of rice and Arabidopsis (Seto *et al*., 2014). It is important to note that in a *Slmax1* TILLING mutant, trace levels of SLs were still detectable in both root exudates and root extracts. However, no germination assay was performed to assess whether these small amounts of SLs were sufficient to induce residual broomrape germination (Zhang *et al*., 2018). It remains unclear if the *Slmax1* lines release additional unknown SL substrates of *SlMAX1* that might trigger the germination of parasitic plant seeds.

The *Slmax1* lines were also characterized by a dramatic reduction in fruit set and by a consistent increase in the time from anthesis to the onset of ripening (breaker); these traits were accompanied by the phenotypes already observed in the *Slccd*s lines (reduction in number of flowers and fruit weight/size). General reproductive defects in the *Slmax1* genotype have been previously reported (Zhang *et al*., 2018), as well as reductions in flower and/or fruit size associated with SL depletion in *Slccd8* lines (Kohlen *et al*., 2012; Bari *et al*., 2019). While the role of SLs in flowering (i.e. induction, number of flowers and inflorescence size) has been extensively studied in tomato and other species, with recent model proposed (Visentin *et al*., 2024 and references therein), the role of SLs in fruit set and ripening remains less understood. In strawberry the expression of the two *MAX1* orthologues was high in style before and after pollination, compared to the lower expression of the *CCD7* orthologue and the negligible expression of the *D27* and *CCD8* orthologues (Wu *et al*., 2019). These findings align with our observations, highlighting a significant contribution of *SlMAX1* to fruit set. This suggests the intriguing possibility of a specific additive effect of the *SlMAX1* loss of function in combination with SL depletion. Additionally, it is plausible that the reproductive and fruit phenotypes observed in *Slccd*s and S*lmax1* lines are linked to the co-regulation of SLs and auxin sensitivity and/or transport (Kohlen *et al*., 2012; Mashiguchi *et al*., 2021). However, the extent of this regulation in fleshy fruits warrants further investigation, and the impact of photosynthate redistribution in highly branched phenotypes cannot be excluded.

### CCDs and MAX1 specifically impact the production of VOCs during fruit ripening

Despite the observed phenotypes in *Slccd*s and *Slmax1* lines, the carotenoid, sugar and flavonoid content in the fruits did not differ significantly from the control. Additionally, water loss, an important determinant of changes in fruit firmness during ripening (Gidado *et al*., 2024), remained unaltered. While these results were not surprising for the *Sld27* lines, given their control-like phenotype, they were unexpected for the lines with more pronounced pleiotropic effects, such as *Slccd*s or *Slmax1*. Interestingly, previous works reported high expression of *CCD7* in tomato fruit at immature green stage (Vogel *et al*., 2010) and of *CCD7* and *CCD8* in young kiwi fruit (*Actinidia chinensis*) (Ledger *et al*., 2010).

To explore potential variations in tomato fruit ripening that may have been overlooked, we analyzed VOCs in fruits from our panel of SL-depleted lines. In addition to their significance as breeding traits for fruit quality, VOCs, deriving from a diversity of metabolic pathways, served as important metabolic checkpoints for possible direct or indirect effects of SLs.

The *Slmax1* fruits lines showed a significant and consistent accumulation of methyl salicylate (MeSA) and guaiacol. MeSA is derived from the methylation of the phytohormone salicylic acid (SA) and acts as a transduction signal in systemic acquired resistance (SAR). In tomato fruit, MeSA concentration typically peak at 15 days after pollination, then decline and remain constant during ripening. As a volatile compound, it is generally associated with a wintergreen flavor, which is often negatively perceived by consumers (Tieman *et al*., 2010; Frick *et al*., 2023; Kaur *et al*., 2023)..

Similarly, guaiacol is synthesized from SA (Mageroy *et al*., 2012; Zhou *et al*., 2021). Guaiacol contributes to a “smoky” or “medicinal” flavor and is prominently emitted in tomato fruits at the mature green stage in both “smoky” and “non-smoky” varieties, although its levels drop dramatically in non-smoky varieties after the breaker stage (Tikunov *et al*., 2013). It is often negatively perceived by consumers (Kaur *et al*., 2023) and the tomato variety used in this work belongs to the “smoky” group, with a high production of guaiacol in the ripe fruit.

In general, changes in in the B pathway can be determined by external causes, such as biotic stress or plant treatments (Huang *et al*., 2020), which could not be excluded in our *Slmax1* lines. However, in Arabidopsis, Kusajima *et al*. (2022) demonstrated that the *max4* mutant (*Slccd8* homologous) exhibited higher free SA levels in leaves compared to the *max3* mutant (the *Slccd7* homologous), which was similar to control levels. This suggests that the perturbation of SA homeostasis may increase with further disruption of the SL biosynthetic pathway.

Amino acid-derived VOCs are key contributors to tomato aroma and flavor, forming during the ripening process (Kaur *et al*., 2023). Consistently with a normal ripening process, the two immediate downstream aldehydes of the amino acid phenylalanine (2-phenylacetaldehyde; floral flavor; Phe) and leucine (3-methylbutanal; malty, earthy flavor; BCAA), both positively contributing to the flavor, were unaltered in SL-depleted lines. In contrast, several downstream VOCs positively contributing to flavor and derived from phenylalanine (2-phenylethanol: nutty, fruity flavor; 1-nitro-2-phenylethane: musty, earthy flavor; Phe) and leucine (1-nitro-3-methylbutane; 3-methylbutanenitrile; BCAA) were dramatically reduced by SL-depletion. Interestingly, we did not observe large variations in the production of VOCs belonging to the ApoC and L pathways, the two more interconnected to carotenoids (Kaur *et al*., 2023).

Therefore, considering the few VOCs altered in the SL-depleted *Sld27* lines, it is likely that the activity of CCD7, CCD8, and MAX1 directly or indirectly influences the ripening-associated biosynthesis of specific VOCs, thereby potentially affecting tomato flavor. The investigation of SLs roles in the ripening processes of both climacteric and non-climacteric species remains an underexplored but compelling area of research (Ferrero *et al*., 2018; Li *et al*., 2023, 2024) and our results contribute valuable insights into these dynamics.

### Field use of SL-depleted tomato lines

Based on morpho-physiological characterization and resistance phenotype, the *D27* gene presents a promising breeding target for developing parasitic plant resistance in tomato. However, the control-like phenotype observed in the *Sld27* lines requires further validation under a broader range of physiological conditions (e.g. abiotic/biotic stress resistance) due to potential overlaps of carotenoid isomerases with other hormonal pathways (*e.g.* abscisic acid) (Tolnai *et al*., 2023; Ye *et al*., 2024). Moreover, field trials are essential to evaluate yield performance comprehensively. The genetic materials developed in this study provide a unique platform for in-depth exploration of SL biosynthesis in a model crop of critical agriculture importance, such as tomato. These resources will be further enriched in the future by incorporating additional CRISPR/Cas9-edited genes involved in the late-step biosynthesis and transport of SLs. The resulting edited plants have the potential to be used not only in field trials as complete specimens but also as rootstock for commercial cultivars/hybrids. Notably, recent field trials have demonstrated the effectiveness of *Slccd7* and *Slccd8* rootstock in controlling *P. aegyptiaca* without yield losses or any apparent pleiotropic effect under normal growing conditions (Karniel *et al*., 2024). However, the use of specific SL-depleted genotypes (*e.g. Slccd7*) as rootstock may also influence physiological traits, such as drought recovery or flowering (Visentin *et al*., 2020, 2024). Additionally, while the use of grafting has shown documented benefits for yield and quality improvement (Caradonia *et al*., 2023; Parisi *et al*., 2023), its adoption in open-field cultivation systems, particularly for processed tomato production, remains constrained due to the higher cost of grafted plants. This underscores the appeal of precise genetic modification (*e.g. D27*) in elite hybrid cultivars as a cost-effective and scalable solution for integrating SL-related resistance traits into commercial production systems.

## Acknowledgments

The authors wish to thank Maurizio Vurro and Angela Boari for their valuable assistance with parasitic weed manipulation. They are grateful to Andrea Mazzucato and Gianfranco Diretto for their guidance on plant phenotypic analysis. Special thanks to Giuseppe Mennella for his advice on metabolic analysis, to Accursio Venezia and Mario Parisi for their support with phytotron and plant management. The authors also appreciate Andrea Burato, Giovanna Forte, Francesco Vitale, and Mario Salzano for their help with plant handling and tomato sampling. Finally, the authors acknowledge Teresa Caballero Vizcaíno and Ana Espinosa Ruiz and the Plant Metabolomics lab of the IBMCP for their technical support in plant metabolomics.

## Author contributions

AN, NDA and TC conceived the experiments. FC, AC and FC assembled the vectors and carried out *in silico* analysis. AN, AC and GF carried out the *in vitro* experiments. AN and AC carried out phenotypic analysis in the phytotron. GF and ADA did the metabolic analysis on sugars and flavonoids. AN, LP and JLR carried out metabolic analysis on carotenoids and VOCs. AN, AC, GF, MMR, JLR and AG analyzed metabolic data and revised results. KT and KY carried out strigolactone analysis on root exudates and performed germination assay. AN, AC, NDA and TC wrote the manuscript. All the authors revised and approved the manuscript.

## Conflict of interest

The authors declare no conflict of interests

## Funding

The work presented here was financially supported by the Italian Ministry of Agriculture, Food Sovereignty and Forests (MASAF), project BIOTECH-Cisget (DM 15924, 18-05-2018).

JLR acknowledges funding from the Spanish Ministry of Science and Innovation by a Juan de la Cierva-incorporación grant (IJC2020-045612-I). AG acknowledges funding from the Spanish Ministry for PGC project PID2022-141438OB-I00 and to the EU for Harnesstom contract 101000716. KY was supported by Japan Science and Technology Agency (FOREST, JPMJFR220F).

## Data availability

The Sanger sequences of target and off-target sites for the *Sld27*, *Slccd7*, *Slccd8* and *Slmax1* lines, including non-edited control plant, are available at Mendeley Data repository (doi: https://doi.org/10.17632/7vwj6cz972.1).

---

**Supplementary Figure S1. Detection of strigolactones in root exudates of non-edited control plants (wild-type, Wt) and CRISPR/Cas9 knock-out (KO) lines targeting core strigolactone biosynthesis genes (*Sld27*, *Slccd7*, *Slccd8* and *Slmax1*)**. Quantification was performed using ultra-performance liquid chromatography coupled with tandem mass spectrometry (UPLC-MS/MS). (A) Multiple reaction monitoring (MRM) chromatograms for orobanchol (m/z 347.1 → 97.0). (B) MRM chromatograms for solanacol (m/z 343.1 → 96.9). (C) MRM chromatograms for 6,7-didehydro-orobanchol, phelipanchol and epiphelipanchol (m/z 345.1 → 97.0). STD: standard

**Supplementary Figure S2**. **Germination-stimulating activity of the tomato root exudates on *Orobanche minor*.** Root exudate samples were collected from both non-edited control plants (wild-type, Wt) and CRISPR/Cas9 knock-out (KO) lines targeting core strigolactone biosynthesis genes (*Sld27*, *Slccd7*, *Slccd8* and *Slmax1*) (for each line, *n* = 3-4). Germination assays were conducted using these root exudates, with GR24 (10[[ M) included as a positive control. Statistically significant differences among genotypes are indicated by different letters (p < 0.05), as determined by Tukey’s HSD test.

**Supplementary Figure S3**. Representative images of stem (A), inflorescence (B), adventitious roots (C) and fruit (D) of the CRISPR/Cas9 edited lines (*Sld27*, *Slccd7*, *Slccd8*, *Slmax1*), compared to non-edited control plants (wild-type, Wt). All images are shown to scale for accurate visual comparison.

## Notes

### Competing Interest Statement

The authors have declared no competing interest.

